# The Pro-vs. Anti-Angiogenic Capacity of Short Chain Fatty Acids is Dependent on the Bioavailability of FFAR3 in Peripheral Artery Disease

**DOI:** 10.1101/2025.02.20.639393

**Authors:** Sivaraman Kuppuswamy, Kripa Patel, Wenbo Zhi, Vijay Ganta

## Abstract

**Background:** Our current study builds on our recent report that showed Palmitate aggravates ischemic endothelial (EC) dysfunction *in vitro* and aims to determine whether Palmitate is a critical determinant of PAD severity. In contrast to Palmitate, we studied the role of short-chain fatty acids (SCFAs) in regulating ischemic revascularization in PAD.

**Methods:** Femoral artery ligation and resection was used as preclinical-PAD model. Hypoxia serum starvation was used as *in vitro* PAD model.

**Results:** Palmitate dramatically decreased ischemic-EC survival and angiogenic capacity *in vitro*, whereas SCFAs significantly induced their angiogenic capacity. LC-MRM MS analysis showed decreased SCFA content in ischemic-muscle. Laser Speckle perfusion imaging showed that intramuscular SCFA treatment significantly induced perfusion recovery, whereas Palmitate showed a modest but significant impairment in perfusion recovery. Immunoblot analysis showed that SCFAs preferentially induce Free fatty acid receptor (FFAR)-3, but not FFAR2. Accordingly, silencing FFAR3 decreased ischemic-EC angiogenic capacity, whereas inhibiting FFAR2 induced ischemic angiogenesis. Pharmacological inhibition of FFAR3 by beta-hydroxybutyrate (BHB) significantly decreased perfusion recovery. SCFA treatment further decreased perfusion recovery in BHB-treated ischemic-muscle. Mechanistically, inhibiting FFAR3-induced FFAR2-levels that inhibited ischemic-EC angiogenic capacity by decreasing NO and inducing ROS levels in SCFA-treated ischemic-ECs.

**Conclusions:** Our data shows that Palmitate alone is not sufficient to drive the PAD severity. Increased FFAR3 levels in ischemic-muscle allow SCFAs to activate the AKT-NO axis to induce ischemic angiogenesis and perfusion recovery. However, loss of FFAR3 in ischemic-muscle promotes FFAR2 activation that blocks SCFA-induced NO-production and redox balance thereby inhibiting perfusion recovery in preclinical-PAD.

## Introduction

Peripheral artery disease (PAD) is a secondary manifestation of atherosclerotic occlusions in the arteries supplying blood to the legs^1^. Even though hemodynamic shear stress at the occluded site favors collateral remodeling by vascular dilation and allows bulk blood flow, progressive atherosclerosis narrows the vascular lumens and blocks the blood flow resulting in impaired microvascular perfusion^2^. In severe PAD (chronic limb-threatening ischemia, CLTI), decreased blood flow and loss of compensatory macro-and microvascular remodeling often result in extensive tissue loss and necrosis leading to approximately 120,000 amputations/year in the US^3,4^. More importantly, the mortality rate in CLTI patients within the first year of diagnosis is ∼25% and a 5-year survival rate is <30%^4^.

Endothelial cells (ECs) are at the center of vascular remodeling processes that drive perfusion recovery in PAD^5^. EC barrier not only limits fluid influx into the tissue but also prevents circulating fatty acids from depositing into the vascular wall^6,7^. Hence, controlling lipid/fatty acid levels is a critical component not only to mitigate atherosclerosis progression but also to reduce the cardiovascular disease burden (CVD)^8^. Fatty acids are sub-grouped into 3 types based on the number of carbon atoms^9^. Short-chain fatty acids (SCFAs) have ≤ 6 carbons, medium-chain fatty acids have 8-12 carbons and long-chain fatty acids (LCFAs) have ≥ 14 carbons. LCFAs are further categorized into saturated and unsaturated fatty acids (monosaturated or polyunsaturated) depending on the carbon double bonds. Saturated fatty acids including Myristic acid, Palmitic acid, Stearic acid, and monounsaturated fatty acids including Palmitoleic acid have been shown to increase the CVD burden^10^, Whereas, monounsaturated Oleic acid and polyunsaturated fatty acids including Linoleic acid, Arachidonic acid, Linolenic acid, Eicosapentaenoic acid, and Docosahexaenoic acid have been shown to decrease CVD burden^11^. Palmitic acid, the most abundant saturated fatty acid has been shown to induce EC-inflammation and contribute to atherosclerosis^12–14^. In fact, a multivariate regression analysis from the Insulin Resistance Atherosclerosis Study^15^ (IRAS) indicated that Palmitic acid is a strong risk factor for Type-2 diabetes (T2D) independent of Insulin resistance.

SCFAs are the metabolic byproducts of dietary fibers (predominantly) by gut bacteria^16^. Acetate>Propionate>Butyrate (60:20:20 mM/kg) are the abundant SCFAs produced in the colon with preferential oxidation towards Butyrate>Propionate>Acetate by gut commensals and intestine for energy production^17^. In a clinical study by Muradi et al^18^ with a small cohort of PAD patients with diabetes, increased fecal excretion of Acetate, Butyrate, and Propionate showed a weak positive correlation with a higher risk of random blood glucose, peak systolic velocity, volume flow, and plaque indicating higher cardiovascular risk in PAD patients with diabetes^18^. Increasing evidence shows that SCFAs improve the atherosclerotic burden^19^, however, the molecular mechanisms by which SCFAs exert their beneficial effects, especially in ECs or in PAD are not known. The mechanism of SCFA action involves signaling through their primary receptors FFAR3 and FFAR2^20^. While Propionate is considered the potent activator of FFAR2, both Propionate, and Butyrate are considered potent activators of FFAR3 followed by Acetate^21,22^. Both FFAR2 and FFAR3 were known to be expressed in skeletal muscle^23^. A 3^rd^ SCFA receptor GPR109a activated by Butyrate was found to be expressed primarily in immune cells and adipocytes^24^.

While SCFAs are produced in the gut, Ketone bodies (e.g. β-hydroxy butyrate (BHB: CH3CH(OH)CH2COO-) structurally similar to the SCFA-Butyrate (CH3CHCH2COO-) are produced in the liver^25^ and extracted into non-hepatic tissues from circulation by SLC16A family members-1 and-7^26^. Despite exhibiting close structural similarities, BHB and Butyrate have been shown to exert distinct functions in ECs^27,28^. More importantly, conflicting reports indicate that while SCFAs function as an agonist for FFAR3, BHB functions as an inhibitor for FFAR3^27,29,30^. Furthermore, while the β-oxidation of LCFAs in mitochondria occurs via the Carnitine-palmitate shuttle system^31^, SCFAs and Ketone bodies can freely diffuse into the mitochondrial matrix and enter the β-oxidation due to their smaller size^32^. Oxidation of Ketone bodies is one of the significant contributors to energy production during starvation/fasting and post-exercise^33,34^. While Ketone bodies also serve in lipogenic and sterol metabolism in the central nervous system and mammary gland, dysfunctional Ketone body metabolism has been shown to occur in multiple diseases including heart failure, aging, and diabetes^25^. In fact, exercise reversed the accumulation of Ketone body (BHB) accumulation and 3-Ketoacid Co-A transferase levels in the gastrocnemius muscle vs. sedentary streptozotocin-induced diabetic rats indicating a negative role of Ketone bodies in Diabetes^35^.

Using *in vitro* T2D-PAD models, we have recently shown a detrimental role of Palmitic acid in aggravating ischemic-EC dysfunction^36^. However, the molecular mechanisms that drive palmitate-induced ischemic-EC dysfunction are not known. Hence, the primary objective of our study was aimed to understand whether Palmitate treatment can impair perfusion recovery in C57BL/6J mice to the same degree observed in T2D-PAD models^36–38^. Based on the emerging roles of SCFAs in regulating cardiovascular diseases^39^, we included SCFA treatment that serves as an additional comparator to Palmitate in determining their roles in regulating ischemic-muscle revascularization and perfusion recovery in PAD.

## Results

### Palmitate induces EC dysfunction, whereas SCFAs improve EC dysfunction

To determine the role of Palmitate in regulating ischemic-EC function, we treated normal or hypoxia serum starved (HSS) immortalized skeletal muscle microvascular ECs (imSkVECs^40^) with Palmitate dose-dependently^36^ and examined EC-functions including proliferation/survival by CCK8, angiogenic-capacity on growth factor reduced Matrigel (GFRM) and barrier-integrity by trans-endothelial electrical resistance experiments. Long-chain fatty acids (LCFAs) and short-chain fatty acids (SCFAs) were used as comparators to Palmitate treatment. While normal-ECs treated with 100µM Palmitate significantly increased EC proliferation (P=0.02), 250µM had no effect, whereas 500µM significantly decreased EC proliferation (P<0.0001, Fig-1A) vs. BSA (vehicle-control for Palmitate) treated controls. LCFAs significantly induced normal-EC survival dose-dependently (1%-P=0.0002, 2.5%-P<0.0001, 5%-P<0.0001) vs. controls (Fig-1A). SCFAs did not affect normal EC proliferation at any concentration. HSS-ECs treated with Palmitate significantly decreased EC survival dose-dependently (100µM:P=0.03, 250µM:P<0.0001, 500µM:P<0.0001, Fig-1B). However, LCFAs or SCFAs did not affect HSS-EC survival vs. control at any doses (Fig-1B).

**Figure 1:**
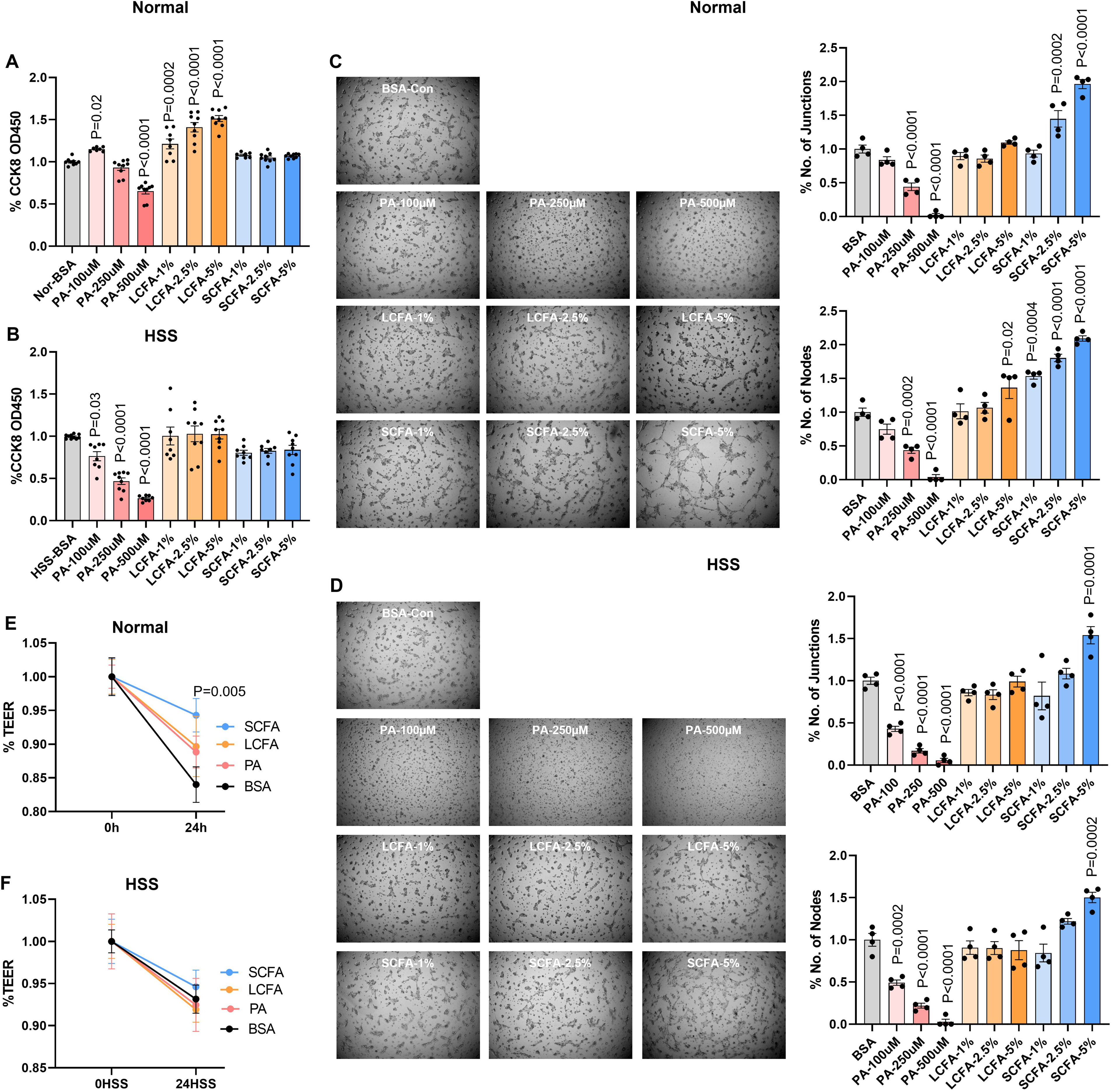
Palmitate, LCFAs and SCFAs have distinct effects on normal vs. ischemic-EC function. A, B) CCK8 cell proliferation/survival assay in A) Normal, B) Hypoxia serum starved (HSS) imSkVECs treated with BSA, Palmitate (PA, 100µM, 250µM, 500µM), long chain fatty acids (LCFAs, 1%, 2.5%, 5%) or short chain fatty acids (SCFAs, 1%, 2.5%, 5%) dose-dependently for 24h. n≥7. One Way ANOVA with Dunnettt’s Post-Test. C, D) *In vitro* angiogenesis assay of C) Normal, D) HSS imSkVECs treated with BSA, Palmitate (100µM, 250µM, 500µM), LCFAs (1%, 2.5%, 5%) or SCFAs (1%, 2.5%, 5%) dose-dependently for 24h on growth factor reduced Matrigel. n=4. One Way ANOVA with Dunnettt’s Post-Test. E, F) Trans-endothelial electrical resistance (TEER, Ohms/cm^2^) of E) Normal, F) HSS imSkVECs treated with BSA, Palmitate (250µM), LCFAs (5%) or SCFAs (5%) for 24h. n=4. Repeated measures ANOVA with Bonferroni select pair comparison. P<0.05 is considered significant.

*In vitro,* angiogenesis assays showed that Palmitate significantly decreased the angiogenic capacity of normal-ECs dose-dependently (250µM:P<0.0001, 500µM:P<0.00011, Fig-1C). While LCFAs did not show any significant effect on the angiogenic capacity of normal-ECs, SCFAs significantly induced normal-EC angiogenic-capacity dose-dependently (%Junctions: 2.5%:P=0.0002, 5%:P<0.0001, %Nodes: 1%:P=0.004, 2.5%:P<0.0001, 5%:P<0.0001 Fig-1C). Palmitate-treated HSS-ECs showed a significant loss of angiogenic capacity dose-dependently (%Junctions 100µM:P<0.0001, 250µM:P<0.0001, 500µM:P<0.0001, %Nodes 100µM:P=0.0002, 250µM:P<0.0001, 500µM:P<0.0001, Fig-1D). While LCFAs did not significantly affect HSS-EC angiogenic capacity, SCFAs significantly induced the angiogenic capacity of HSS-ECs (%Junctions 5%:P=0.0001, %Nodes: 5%:P=0.0002, Fig-1D).

Since Palmitate showed a detrimental effect on normal and HSS-EC survival and angiogenic capacity at the highest 500µM concentration, we used 250µM Palmitate concentration for measuring EC barrier integrity). ECs treated with Palmitate or LCFAs did not show a significant effect on the EC barrier under normal growth conditions. However, consistent with the role of SCFA in promoting the epithelial barrier^41^, normal-ECs treated with SCFA showed a significant increase in barrier integrity (P=0.005, Fig-1E). However, HSS-ECs treated with Palmitate, LCFAs, or SCFAs showed no significant effect on barrier integrity vs. vehicle control (Fig-1F). These data showed selective vascular effects of Palmitate, LCFAs, and SCFA in regulating normal and ischemic-EC functions.

### SCFAs induce distinct metabolic signatures from Palmitate and LCFAs in normal and HSS-ECs

Since normal-ECs use glycolysis as the primary pathway for metabolic requirement^42^, we first examined the effect of Palmitate, LCFAs and SCFAs in regulating normal-EC glycolysis. Seahorse-glycolysis stress assay showed no significant effect of Palmitate on normal-EC glycolysis. However, both LCFAs and SCFAs significantly increased glycolysis (LCFA:P<0.0001, SCFA-P=0.004), glycolytic-capacity (LCFA:P<0.0001, SCFA:P<0.0001) and glycolytic-reserve (LCFA:P<0.0001, SCFA:P<0.0001) vs. vehicle-control (Fig-2A). In HSS-ECs, Palmitate treatment significantly decreased glycolysis (P<0.0001), glycolytic-capacity (P<0.0001), and glycolytic-reserve (P<0.0001) vs. vehicle-control (Fig-2B). LCFAs significantly induced glycolysis (P=0.02) but did not affect glycolytic-capacity or glycolytic-reserve vs. control (Fig-2B). SCFAs showed a modest but significant decrease in glycolysis (P=0.008), glycolytic-capacity (P=0.01), and glycolytic-reserve (P=0.002) vs. controls (Fig-2B).

**Figure 2:**
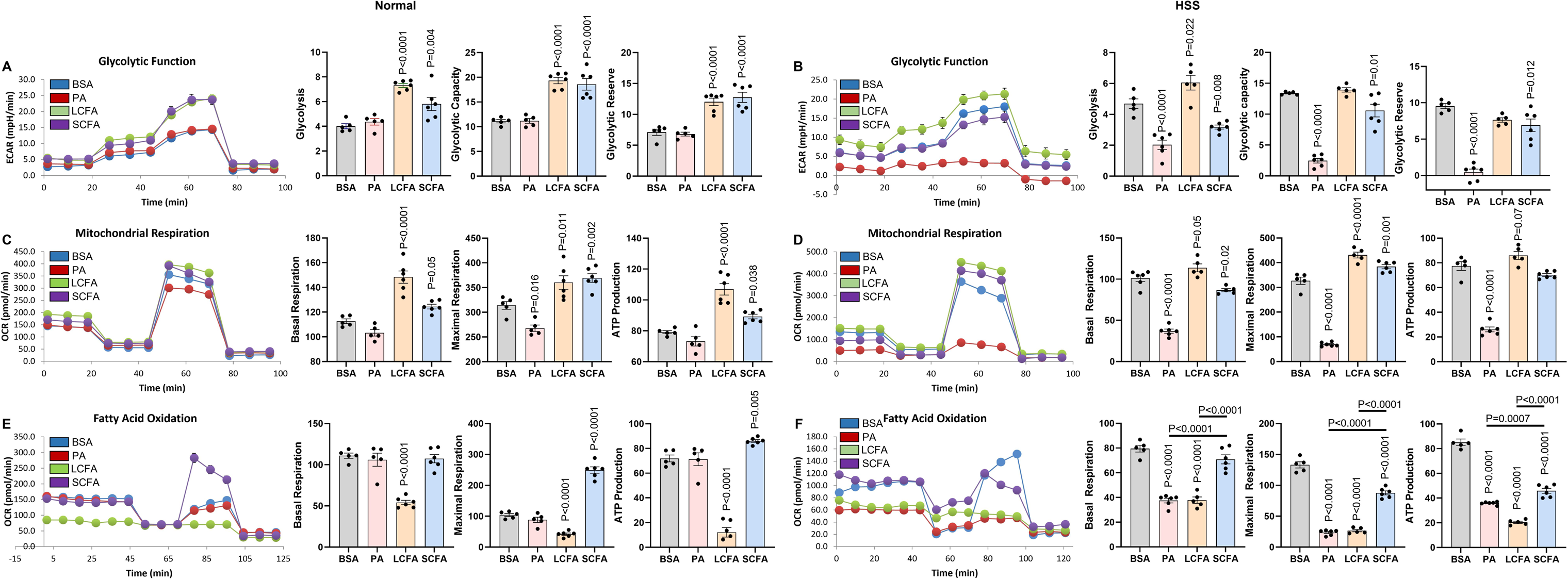
Metabolic regulation of Palmitate, LCFAs, and SCFAs of normal vs. ischemic-EC. A, B) Seahorse Glycolysis stress test in A) Normal, B) Hypoxia serum starved (HSS) imSkVECs treated with BSA (n=5), Palmitate (PA, 250µM, n=5), long chain fatty acids (LCFA, 5%, n=6), or short chain fatty acids (SCFAs, 5%, n=6) for 24h. One Way ANOVA with Dunnettt’s Post-Test. C, D) Seahorse Mitostress test in A) Normal, B) HSS imSkVECs treated with BSA (n=5), Palmitate (PA, 250µM, n=6), long chain fatty acids (LCFA, 5%, n=5), or short chain fatty acids (SCFAs, 5%, n=6) for 24h. One Way ANOVA with Dunnettt’s Post-Test. E, F) Seahorse basal Fatty acid Oxidation test (that uses CPT1 inhibitor without exogenous/additional Palmitate in the Seahorse assay medium) in A) Normal, B) HSS imSkVECs treated with BSA (n=5), Palmitate (PA, 250µM, n=6), long chain fatty acids (LCFA, 5%, n=5), or short chain fatty acids (SCFAs, 5%, n=6) for 24h. One Way ANOVA with Dunnettt’s Post-Test. P<0.05 is considered significant.

While impaired glycolysis explains the loss of angiogenic capacity in Palmitate-treated HSS-ECs, decreased glycolysis in SCFA-treated HSS-ECs didn’t explain the ability of SCFA to induce HSS-EC angiogenic capacity. Since fatty acids undergo β-oxidation in mitochondria^31^, we next examined the role of Palmitate, LCFAs, and SCFAs in regulating mitochondrial respiration/OxPhos. Seahorse Mitostress assay showed that Palmitate significantly decreased maximal-respiration (P=0.016) but not basal-respiration or ATP-production in normal-ECs vs. vehicle-control (Fig-1C). Both LCFAs and SCFAs significantly increased basal-respiration (LCFAs:P<0.0001, SCFAs:P=0.05), maximal-respiration (LCFAs:P<0.011, SCFAs:P=0.002), and ATP-production (LCFAs:P<0.0001, SCFAs:P=0.038) vs. vehicle-controls (Fig-2C). In HSS-ECs, Palmitate significantly decreased basal-respiration (P<0.0001), maximal-respiration (P<0.0001), and ATP-production (P<0.0001) vs. control (Fig-2D). LCFAs significantly induced basal-respiration (P=0.05) and maximal-respiration (P<0.0001) with a numerical increase in ATP-production (P=0.07) vs. control (Fig-2D). SCFAs significantly decreased basal-respiration (P=0.02) but induced maximal-respiration (P=0.001) in HSS-ECs with no changes in ATP-production vs. control (Fig-2D).

While LCFAs use the carnitine-shuttle system for β-oxidation in mitochondria^31^, SCFAs can directly enter (by diffusion) mitochondria for β-oxidation^32^. Hence, we next determined the role of fatty acid oxidation (FAO) in regulating normal and HSS-EC metabolic/functional responses to Palmitate, LCFA, and SCFA. Since, ECs are already treated with Palmitate, LCFAs, and SCFAs, Seahorse basal-FAO assay that uses CPT1 inhibitor, Etoxomir to block Carnitine-shuttling into mitochondria in the absence of additional Palmitate in the seahorse basal medium was used. FAO assay showed no significant effect of Palmitate or SCFAs in regulating normal-EC basal-respiration. However, normal-ECs treated with LCFAs showed a significant loss of basal-respiration (P<0.0001), maximal-respiration (P<0.0001), and ATP-production (P<0.0001) vs. control (Fig-2E). While Palmitate did not affect mitochondrial-respiration, SCFAs treated normal-ECs showed a significant increase in maximal-respiration (P<0.0001) and ATP-production (P<0.0001) vs. control. On the contrary, HSS-ECs treated with Palmitate, and LCFAs showed a significant loss of basal-respiration (P<0.0001), maximal-respiration (P<0.0001), and ATP-production (P<0.0001) vs. control (Fig-2F). SCFAs treated HSS-ECs showed no significant difference in basal-respiration but showed a significant decrease in maximal-respiration (P<0.0001) and ATP-production (P<0.0001) vs. controls (Fig-2F). However, vs. Palmitate and LCFAs treated HSS-ECs, SCFAs treated HSS-ECs showed a significant increase in basal-respiration and maximal-respiration (P<0.0001 vs. Palmitate or LCFAs), and ATP-production (P=0.0007 vs. Palmitate, (P<0.0001 vs. LCFA, Fig-2F). This data indicated that the ability of Palmitate and LCFAs to exert a metabolic effect in HSS-ECs is strictly dependent on the carnitine-shuttle system (regulated by CPTs) but not for SCFA. Furthermore, no changes in ATP production by SCFAs suggested that the ability of SCFAs to induce ischemic-EC angiogenic capacity is independent of additional ATP-production.

### Palmitate inhibits perfusion recovery whereas SCFAs induce perfusion recovery

To determine the pathological relevance of increased Palmitate levels in inhibiting perfusion recovery in PAD, we treated C57BL/6J mice with Palmitate (pH balanced 100µl of 250µM BSA-Palmitate i.m. at 3 sites in skeletal muscle (2 non-overlapping sites in GA and 1 site in TA (∼30ul solution/site)). Control mice received BSA (v/v, i.m.) at d0, d3, d7, d14 and d21 post HLI (Fig-3A). Since SCFAs significantly induced the angiogenic capacity of ischemic-ECs, we also wanted to determine the functional role of SCFAs in preclinical-PAD. However, limited information is available on the levels or function of SCFAs in PAD. Hence, we performed LC-MSMS analysis to determine SCFA levels in normal vs. ischemic-muscle and serum samples at an informative day-3 post-HLI^36,37,40,43–45^. LC-MSMS analysis showed no significant difference in the levels of Propionate, Butyrate, Hexanoic, Heptanoic, Octanoic, Nonanoic, and Decanoic acid levels between normal vs. ischemic-muscle samples (Fig-3A). However, a significant decrease (P<0.00069) in Dodecanoic acid levels is observed in ischemic-muscle vs. normal muscle (Fig-3A). No significant difference in the circulating levels of Propionate, Butyrate, Hexanoic, Heptanoic, Octanoic, Nonanoic, Decanoic, or Dodecanoic acid was observed between ischemic vs. normal serum samples (Fig-3B). This data indicated that HLI neither increases the circulating SCFA levels nor their uptake into the ischemic-muscle.

**Figure 3:**
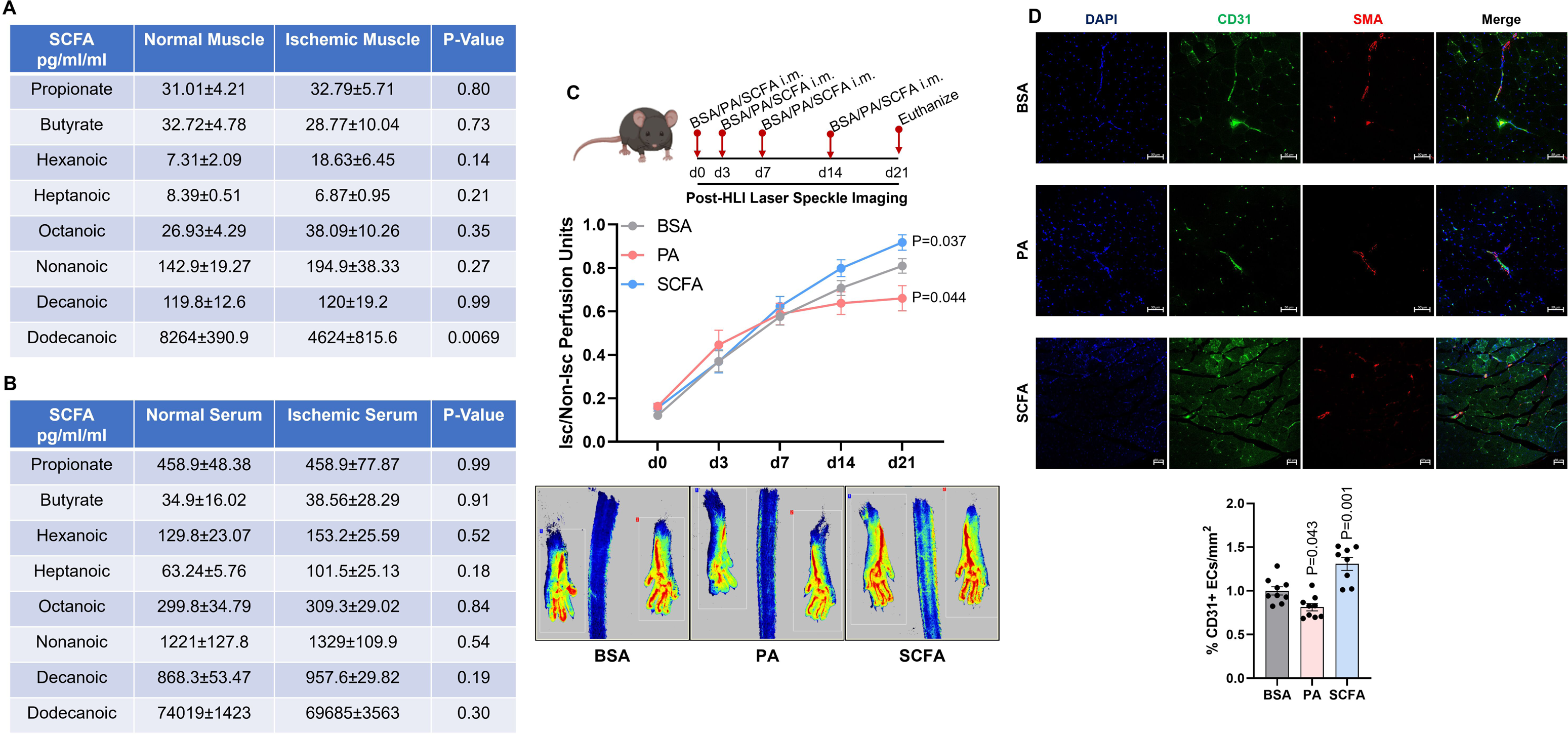
Palmitate inhibits, whereas SCFAs induce Perfusion recovery in experimental-PAD. A, B) LC-MSMS analysis of SCFA levels in A) normal (n=4) vs. ischemic-muscle (n=4) and B) serum from normal (n=4) and ischemic mice (n=4) at day-3 post HLI. Unpaired T-Test between specific SCFAs. C) Laser speckle perfusion imaging in C57BL/6J mice treated with BSA (n=10), Palmitate (PA, 250µM, n=9) or SCFAs (500µM, n=12) at d0, d3, d7 and d14 post HLI i.m. Repeated measures ANOVA with Bonferroni select pair comparison. D) Immunohistochemistry of CD31 and α-SMA in the ischemic-muscle of mice treated with BSA (n=9), Palmitate (PA, 250µM, n=9) or SCFAs (500µM, n=8) at d21 post-HLI. One Way ANOVA with Dunnettt’s Post-Test. P<0.05 is considered significant.

Laser Speckle perfusion imaging showed that while Palmitate significantly decreased the perfusion recovery in C57BL/6J mice (d21 post-HLI: BSA-0.810±0.33 vs. PA-0.661±0.058, P=0.037, Fig-3B), mice that received SCFAs (pH balanced 100ul of 500µM SCFAs i.m. at 3 sites in skeletal muscle (2 non-overlapping sites in GA and 1 site in TA (∼30µl solution/site)) in parallel to Palmitate showed a significant increase in perfusion recovery vs. vehicle-controls (Fig-3B). In preclinical-PAD models, C57BL/6J mice strain shows excellent perfusion recovery^45^. Hence it is often difficult to observe an enhanced perfusion recovery with therapeutics that can promote perfusion recovery in this mouse strain. However, C57BL/6J mice treated with SCFAs showed a modest but significant increase in perfusion recovery vs. vehicle-control (d21 post-HLI: BSA-0.810±0.033 vs. SCFA-0.918±0.035, P=0.044). Taking the mouse stain into consideration, the ability of SCFAs to further induce perfusion recovery in this mouse strain indicates a profound effect of SCFAs’ ability to induce perfusion recovery in PAD. Since LCFAs did not show any significant effect on EC function in vitro, LCFA treatment was not included in the perfusion recovery experiments.

Consistent with the loss of perfusion recovery, microvascular density assessed by CD31+ECs/mm2 and macrovascular density assessed by SMA+ vessels >25um/mm2 showed that Palmitate significantly decreased microvascular density (P=0.021), whereas SCFAs significantly induced (P=0.001) microvascular density in the ischemic-muscle vs. vehicle-control (Fig-3D). No significant differences in the macrovascular density were observed among the groups. These data indicated the therapeutic potential of SCFAs in driving perfusion recovery in PAD.

### FFAR3 bioavailability regulates the ability of SCFAs to revascularize ischemic-muscle in preclinical-PAD

We next wanted to determine the mechanism by which SCFAs regulate ischemic-EC angiogenic capacity and perfusion recovery is preclinical-PAD. Since FFAR2 and FFAR3 are the principal receptors for SCFAs^23^, we examined whether SCFAs preferentially induce FFAR2 or FFAR3 levels in ischemic-ECs. Western blot analysis showed that while Palmitate (P=0.022) and LCFAs (P=0.003) significantly induce FFAR2 levels in ischemic-ECs, whereas SCFAs did not show a significant effect on the FFAR2 levels vs. controls (Fig-4A). On the contrary, while Palmitate or LCFAs did not affect FFAR3 levels in ischemic-ECs, SCFAs significantly induced FFAR3 levels (P<0.0001) in ischemic-ECs vs. control (Fig-4A). A comparison of non-ischemic vs. ischemic-ECs showed that ischemia induces FFAR3 levels but not FFAR2 levels (Fig-4B) While a significant increase in both FFAR2 and FFAR3 levels was observed in the ischemic-muscle at an informative day-3 post-HLI, the magnitude of FFAR3 expression (∼2X) was higher than FFAR2 expression (0.7X) in the ischemic vs. non-ischemic-muscle (Fig-4C). Consistently, immunohistochemical analysis to determine EC-specific FFAR3 expression showed a significant increase in the numbers of CD31^+^FFAR3^+^ ECs in the ischemic-muscle vs. non-ischemic (Fig-4D). These data indicated that ischemia-induced FFAR3 levels increase their bioavailability to exogenously delivered SCFAs to promote perfusion recovery in preclinical-PAD.

**Figure 4:**
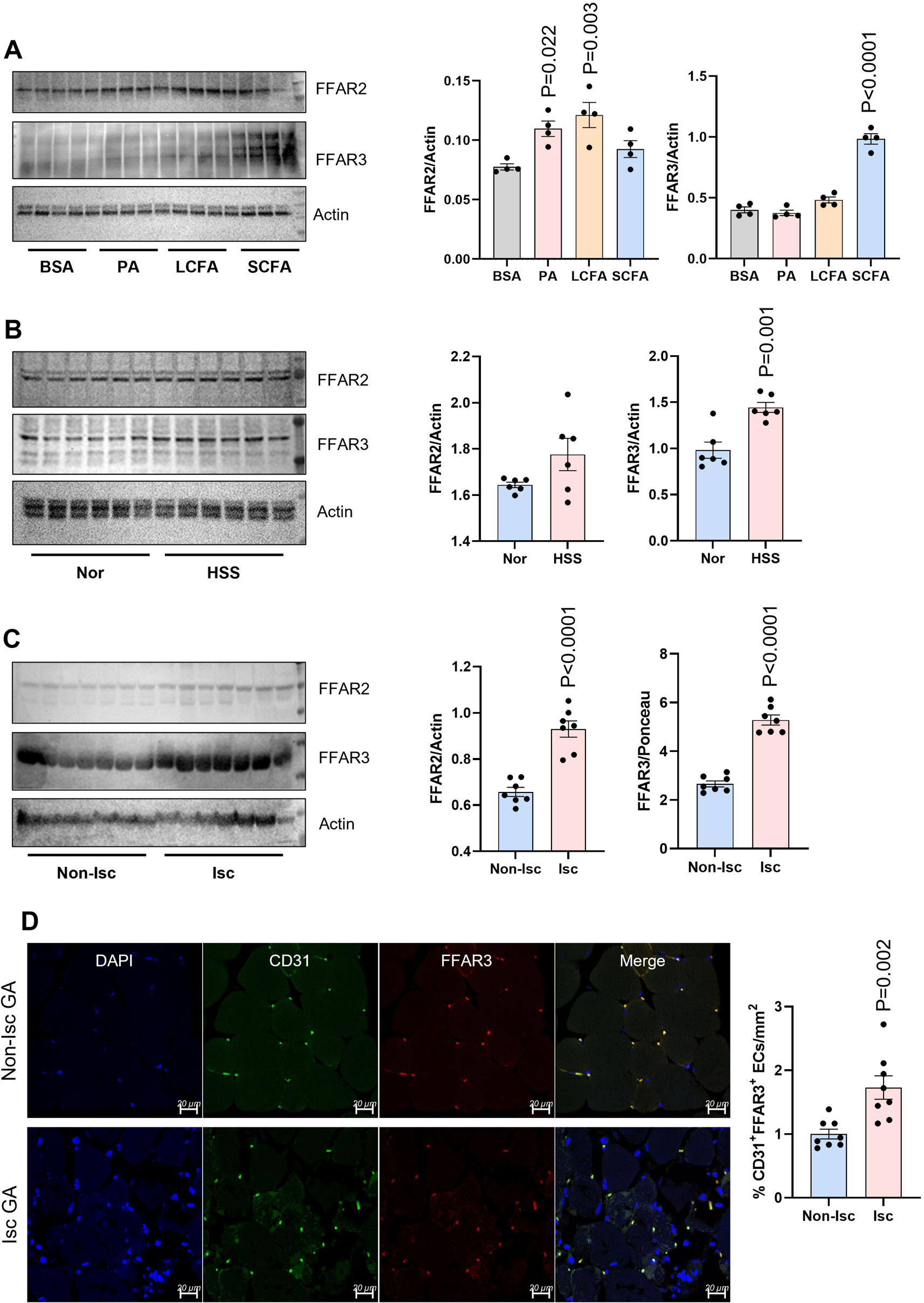
Ischemia induces FFAR3 levels in ischemic-muscle and HSS ECs. A) Western blot analysis of FFAR2 and FFAR3 in HSS imSkVECs treated with BSA (n=4), PA (250uM, 24h), LCFA (5%, 24h) or SCFA (5%, n=4) for 24h. One Way ANOVA with Dunnettt’s Post-Test. B) Western blot analysis of FFAR2 and FFAR3 in normal (n=6) vs. 24h HSS imSkVECs (n=6). Unpaired T-test. C) Western blot analysis of FFAR2 and FFAR3 in non-ischemic (n=7) vs. ischemic (n=7) gastrocnemius muscle at day-3 post HLI. Unpaired T-test. C) Immunohistochemical analysis of CD31 and FFAR3 in non-ischemic (n=8) vs. ischemic (n=8) gastrocnemius muscle tissue sections at day-7 post HLI. Unpaired T-test. P<0.05 is considered significant.

Next, to confirm the causal role of FFAR3 in driving the angiogenic capacity and perfusion recovery induced by SCFAs, we performed FFAR3 loss of function experiments in the presence of SCFAs. Even though existing literature is conflicting, to date, β-hydroxy butyrate (BHB) is the only known inhibitor of FFAR3 and HDAC1-3^27,46^. Hence, we first confirmed the specific inhibitory property of BHB towards HDACs vs. FFAR3 in ischemic-ECs. Western blot analysis showed ischemic-ECs treated with BHB showed no significant difference in the HDAC1, HDAC2, or HDAC3 levels, but a significant decrease in FFAR3 levels vs. controls (Fig-5A). Consistent with the BHB action, silencing FFAR3 also did not show any significant difference in HDAC1, HDAC2, or HDAC3 levels vs. control-siRNA in ischemic-ECs (Fig-5B). To determine whether the perfusion recovery induced by SCFAs is dependent on FFAR3, we pre-treated mice with BHB (pH balanced, 5mM, i.m. 100ul, 3 sites, 2 sites in GA and 1 site in TA) a day before HLI. The following day, mice were treated with HBSS (100µl), BHB (pH balanced, 5mM) or BHB+SCFA (pH balanced, 5mM BHB+500µM SCFA) i.m immediately post HLI (d0) and at day-3, 7, and 14. We anticipated that BHB-treated mice would show an impaired perfusion recovery and SCFA treatment would fail to improve perfusion recovery in BHB-treated mice. However, while BHB treatment showed a modest but significant decrease in perfusion recovery vs. control at day21 post-HLI (HBSS:0.828±0.032 vs. BHB:0.719±0.043), mice that received a combination of BHB+SCFA treatment showed a further loss of perfusion recovery (HBSS:0.828±0.032 vs. BHB+SCFA: 0.595±0.035, Fig-5C). More importantly, mice treated with BHB+SCFA showed a numerical increase in necrosis incidence vs. BHB or control mice (Fig-5D). Immunohistochemical analysis of CD31 and SMA showed a significant decrease (P=0.001) in CD31^+^ ECs in ischemic-muscle treated with BHB vs. control. SCFA treatment further decreased (P<0.0001) CD31^+^ ECs in the ischemic-muscle vs. BHB or control mice (Fig-5E). No significant difference in arteriolar density was observed among the groups. These data indicated that loss of FFAR3 bioavailability to SCFAs shifts their pro-angiogenic property to anti-angiogenic in the ischemic-muscle.

**Figure 5:**
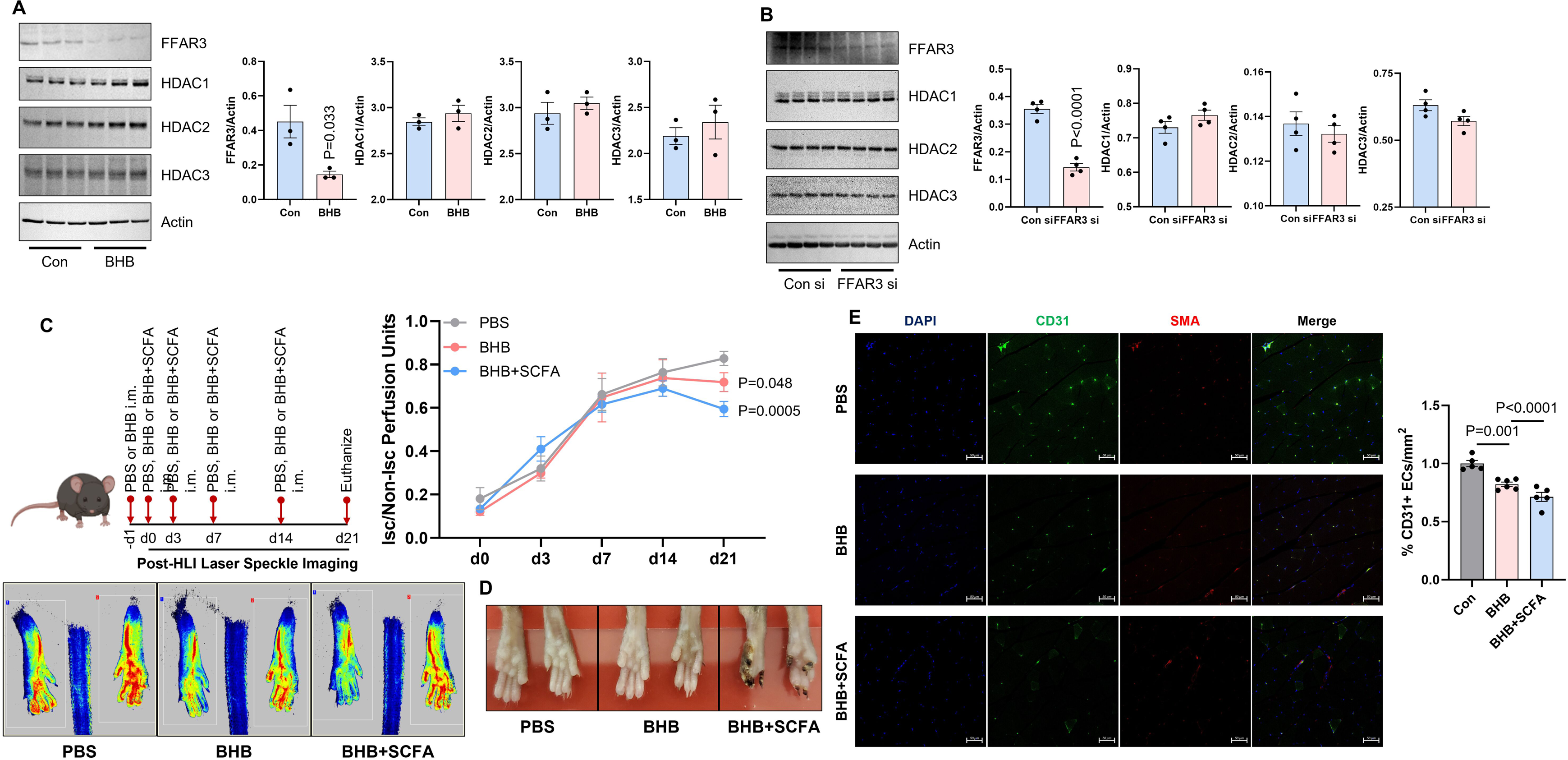
SCFAs impair perfusion recovery upon FFAR3 inhibition in experimental-PAD. A, B) Western blot analysis of FFAR3, HDAC1, HDAC2, HDAC3, Actin in HSS imSkVECs treated with A) Con (n=3) or BHB (5mM, n=3), B) Con si (n=4) or FFAR3 si (n=4). Unpaired T-Test. C) Laser speckle perfusion imaging in C57BL/6J mice treated with Control (n=6), β-hydroxy butyrate (BHB, 5mm, n=7) or SCFA (500µM) + BHB (5mM) n=7 at-d1, d0, d3, d7 and d14 post HLI i.m. Repeated measures ANOVA with Bonferroni select pair comparison. D) Images of necrotic toes in HLI mice treated with BSA, BHB or BHB+SCFA. E) Immunohistochemistry of CD31 and α-SMA in the ischemic-muscle of mice treated with Control (n=), BHB (n=) or BHB+SCFA (n=8) at d21 post-HLI. One Way ANOVA with Dunnettt’s Post-Test. P<0.05 is considered significant.

Consistent with the impaired perfusion recovery in BHB+SCFA-treated ischemic-muscle, *in vitro* ischemic-ECs treated with BHB showed a dose-dependent decline in survival vs. control. While 1mM BHB significantly increased ischemic-EC survival (P=0.0003), 2.5mm BHB did not have any significant effect and 5mM BHB significantly decreased ischemic-EC survival (P<0.0001, Fig-6A). Ischemic-ECs treated with a combination of BHB+SCFA showed a significant decrease in cell survival vs. control or BHB-treated ischemic-ECs (P=0.0004, Fig-6A). *In vitro* angiogenesis assays showed that while BHB-treated ischemic-ECs showed a loss of angiogenic capacity (% No. of Nodes: P=0.005, % no. of Junctions: P=0.001 and % No. of Branches: P=0.003), BHB+SCFA treated ischemic-ECs showed a further loss of angiogenic-capacity vs. control or BHB-treated ischemic-ECs (% No. of Nodes: P<0.001, % No. of Junctions: P=0.064, % No. of branches: P=0.021 vs. BHB, Fig-6B). Seahorse Mitostress assay showed no significant differences in the basal-respiration but a significant decrease in the maximal respiration of ischemic-ECs treated with BHB vs. control (P=0.035, Fig-6C). BHB+SCFA further decreased the maximal respiratory capacity of ischemic-ECs vs. control or BHB treatments (P=0.001 vs. BHB, Fig-6C). Silencing FFAR3 recapitulated the effects of BHB on ischemic-ECs. CCK8 assay showed that silencing FFAR3 significantly decreased ischemic-EC survival vs. control-siRNA (P=0.003, Fig-6D). SCFA treatment further decreased the survival of FFAR3 silenced ischemic-ECs (P=0.04 vs. FFAR3-si, Fig-6D). *In vitro* angiogenesis assay showed that silencing FFAR3 decreased the angiogenic capacity of ischemic-ECs (% No. of Nodes: P=0.011, % No. of Junctions: P=0.001, % No. of Branches: P=0.008, Fig-6E), that was further decreased by SCFAs vs. FFAR3-si or control-si (% No. of Nodes: P=0.0008, % No. of Junctions: P=0.0001, % No. of Branches: P=0.014 vs. FFAR3-si, Fig-6E). Seahorse Mitostress assay showed that FFAR3-si did not affect basal-respiration but decreased the maximal-respiration of ischemic-ECs (P=0.01) that was further decreased by SCFAs vs. FFAR3-si or control-si (P=0.064 vs. FFAR3-si, Fig-6F). No significant differences in ATP-production were observed among any of the BHB or FFAR3-si treated groups (Fig-6C, F). These data indicated that loss of FFAR3 function shifts the pro-angiogenic capacity of SCFAs to inhibitory in ischemic-ECs independent of ATP-production.

**Figure 6:**
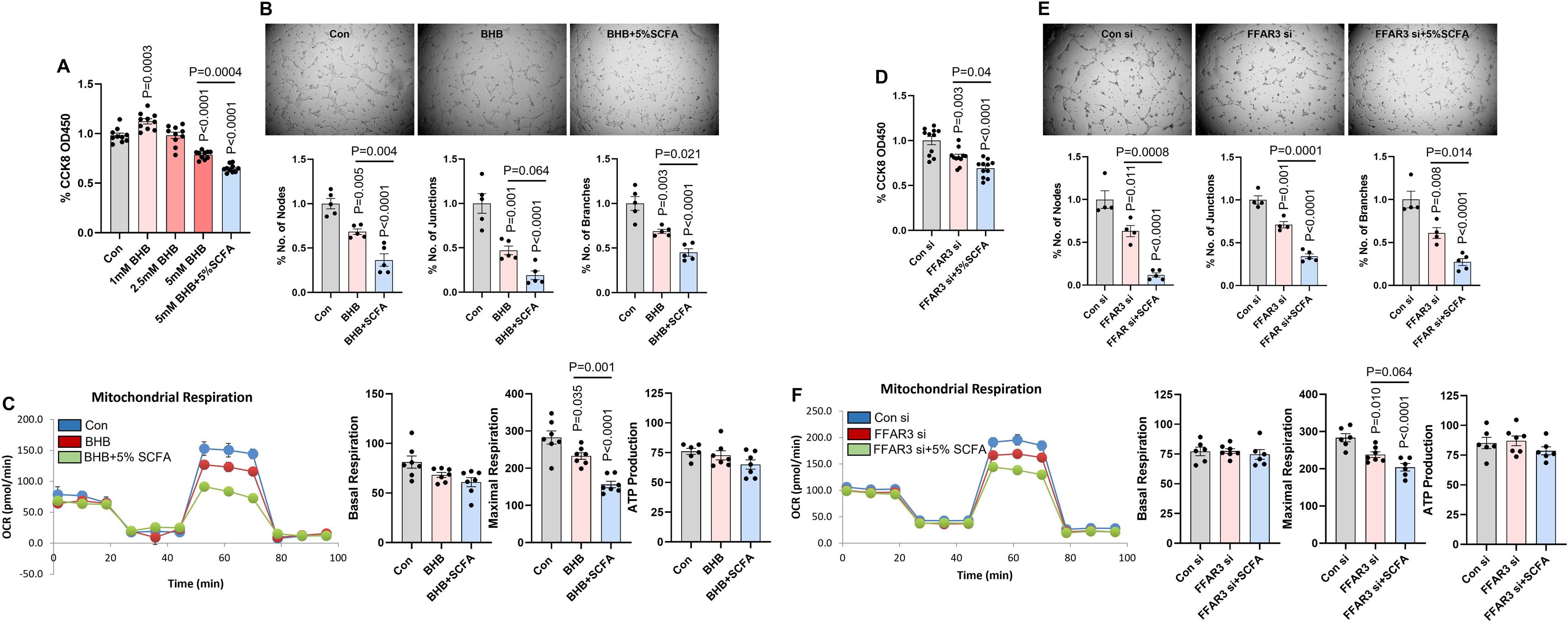
FFAR3 inhibition reverses pro-angiogenic SCFAs to Anti-angiogenic in ischemic-ECs. A, D) CCK8 analysis of HSS imSkVECs treated with A) control, β-hydroxy butyrate (BHB, 1mM, 2.5mM, or 5mM) or BHB (5mm)+SCFA (500µM) for 24h. n=10. One Way ANOVA with Bonferroni select pair comparison B) Con si, FFAR3 si or FFAR3 si+SCFA (500µM) for 24h. n=11. One Way ANOVA with Bonferroni select pair comparison. B, E) *In vitro* angiogenesis assay of HSS imSkVECs treated with B) control, BHB (5mM) or BHB (5mM)+SCFA (500µM) for 24h. n=5. One Way ANOVA with Bonferroni select pair comparison. E) Con si, FFAR3 si or FFAR3 si+SCFA (500µM) for 24h. n≥4. One Way ANOVA with Bonferroni select pair comparison. C, F) Seahorse Mitostress assay in HSS imSkVECs treated with C) control, BHB (5mM) or BHB (5mM)+SCFA (500µM) for 24h. n=7. One Way ANOVA with Bonferroni select pair comparison. F) Con si, FFAR3 si or FFAR3 si+SCFA (500µM) for 24h. n≥6. One Way ANOVA with Bonferroni select pair comparison. P<0.05 is considered significant.

### FFAR3 inhibition induces SCFA-FFAR2 mediated anti-angiogenic properties in ischemic-ECs

We next wanted to determine the mechanism by which SCFAs impaired perfusion recovery upon FFAR3 inhibition. Based on the recent report by Nooromid et al^47^, we reasoned whether inhibiting FFAR3 in the ischemic-ECs allowed the activation of FFAR2 by SCFAs resulting in impaired ischemic-angiogenesis. Western blot analysis showed that silencing FFAR3 (Fig-7A) or treatment with BHB (Fig-7B) significantly increased FFAR2 levels (Con vs. BHB: P=0.001, Fig-7A; Con-si vs. FFAR3-si: P=0.025 Fig-7B) in ischemic-ECs vs. respective controls. While the magnitude of FFAR2 induction achieved by BHB treatment did not allow further increase by SCFAs (Con vs. BHB+SCFA, P=0.009, Fig-7A), SCFA treatment further increased FFAR2 levels in FFAR3 silenced ischemic-ECs (FFAR3-si vs. FFAR3-si+SCFA: P=0.004, Fig-7B). Ischemic-ECs treated with SCFAs dose-dependently did not show any significant effect on HDAC1-3 levels in (Fig-S3). These data indicated that FFAR3 inhibition induces FFAR2 levels that were further induced by SCFA treatment in ischemic-ECs.

**Figure 7:**
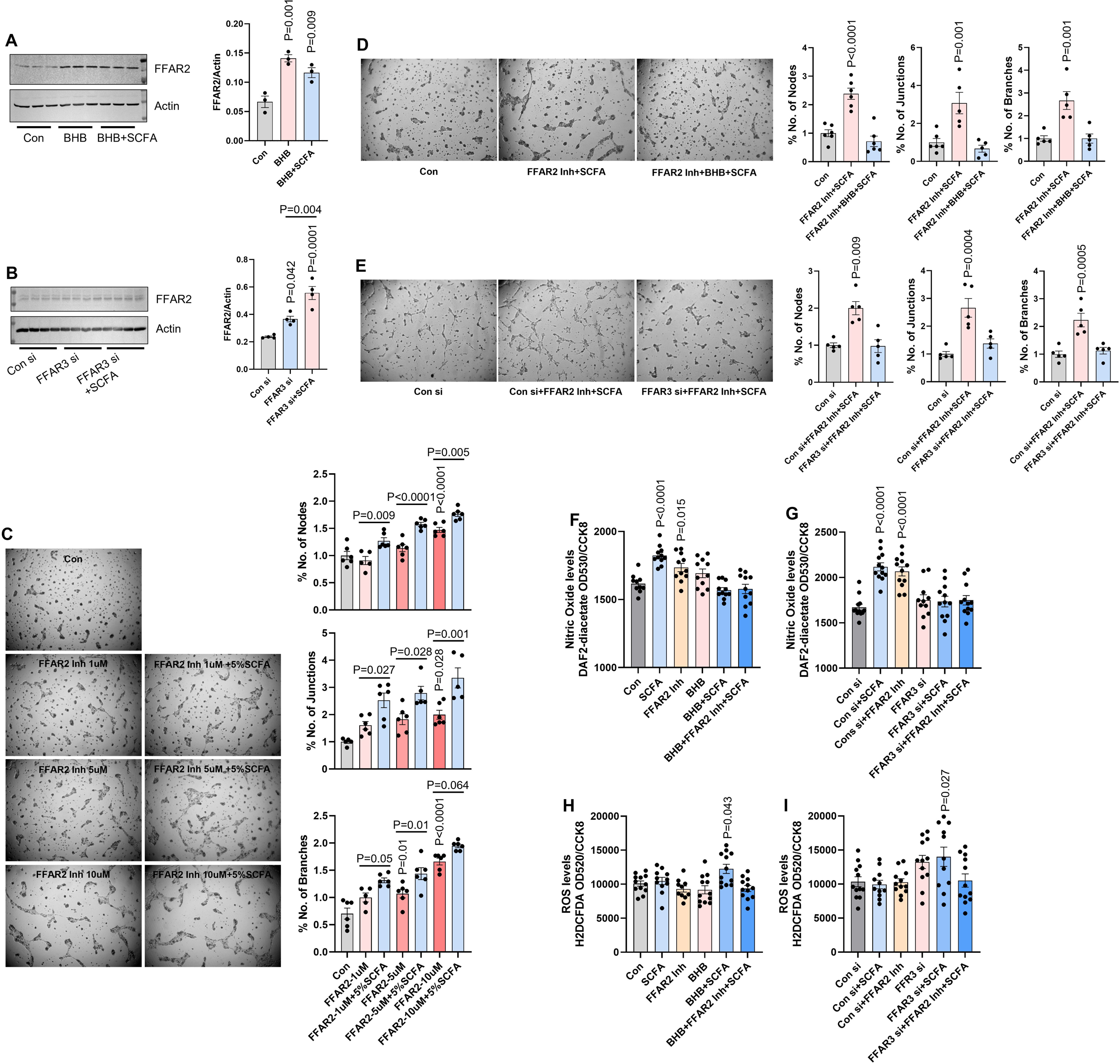
FFAR3 inhibition induces FFAR2 that blocks SCFA-induced ischemic angiogenesis. A, B) Western blot analysis of FFAR2 in HSS imSkVECs treated with A) control, BHB (5mM) or BHB (5mM)+SCFA(500µM) for 24h. n=3. One Way ANOVA with Dunnettt’s Post-Test. B) control si, FFAR3 si or FFAR3 si+SCFA (500µM). n=4. One Way ANOVA with Bonferroni select pair comparison. C, D, E) *In vitro* angiogenesis assay of HSS imSkVECs treated with C) FFAR2 inhibitor (1µM, 5µM, or 10µM) with or without SCFA (500µM) for 24h. One Way ANOVA with Bonferroni select pair comparison. n≥6. D) Control, FFAR2 Inh (10µM)+SCFA (500µM) or FFAR2 Inh (10µM)+BHB (5mM)+SCFA (500µM). n≥6. One Way ANOVA with Dunnettt’s Post-Test. E) Con si, FFAR2 Inh (10µM)+SCFA (500µM), FFAR2 Inh (10µM)+FFAR3 si+SCFA (500µM). n=5. One Way ANOVA with Dunnettt’s Post-Test. F, G) Intracellular Nitric Oxide assay by cell permeable fluorescent DAF2-acetate normalized to CCK8 in HSS imSkVECs treated with F) control, SCFA (500µM), FFAR2 Inh (10µM), BHB (5mM), BHB (5mM)+SCFA (500µM) or BHB (5mM)+FFAR2 Inh (10µM)+SCFA (500µM) for 24h. n=12. One Way ANOVA with Dunnettt’s Post-Test. G) control si, control si+SCFA (500µM), control si+FFAR2 Inh (10µM), FFAR3 si, FFAR3 si+SCFA (500µM) or FFAR3 si+FFAR2 Inh (10µM)+SCFA (500µM) for 24h. n=12. One Way ANOVA with Dunnettt’s Post-Test. H, I) Intracellular ROS levels by cell permeable fluorescent H2DCFDCA normalized to CCK8 in HSS imSkVECs treated with H) control, SCFA (500µM), FFAR2 Inh (10µM), BHB (5mM), BHB (5mM)+SCFA (500µM) or BHB (5mM)+FFAR2 Inh (10µM)+SCFA (500µM) for 24h. n≥10. One Way ANOVA with Dunnettt’s Post-Test. I) control si, control si+SCFA (500µM), control si+FFAR2 Inh (10µM), FFAR3 si, FFAR3 si+SCFA (500µM) or FFAR3 si+FFAR2 Inh (10µM)+SCFA (500µM) for 24h. n=11. One Way ANOVA with Dunnettt’s Post-Test. P<0.05 is considered significant.

**Figure 8:**
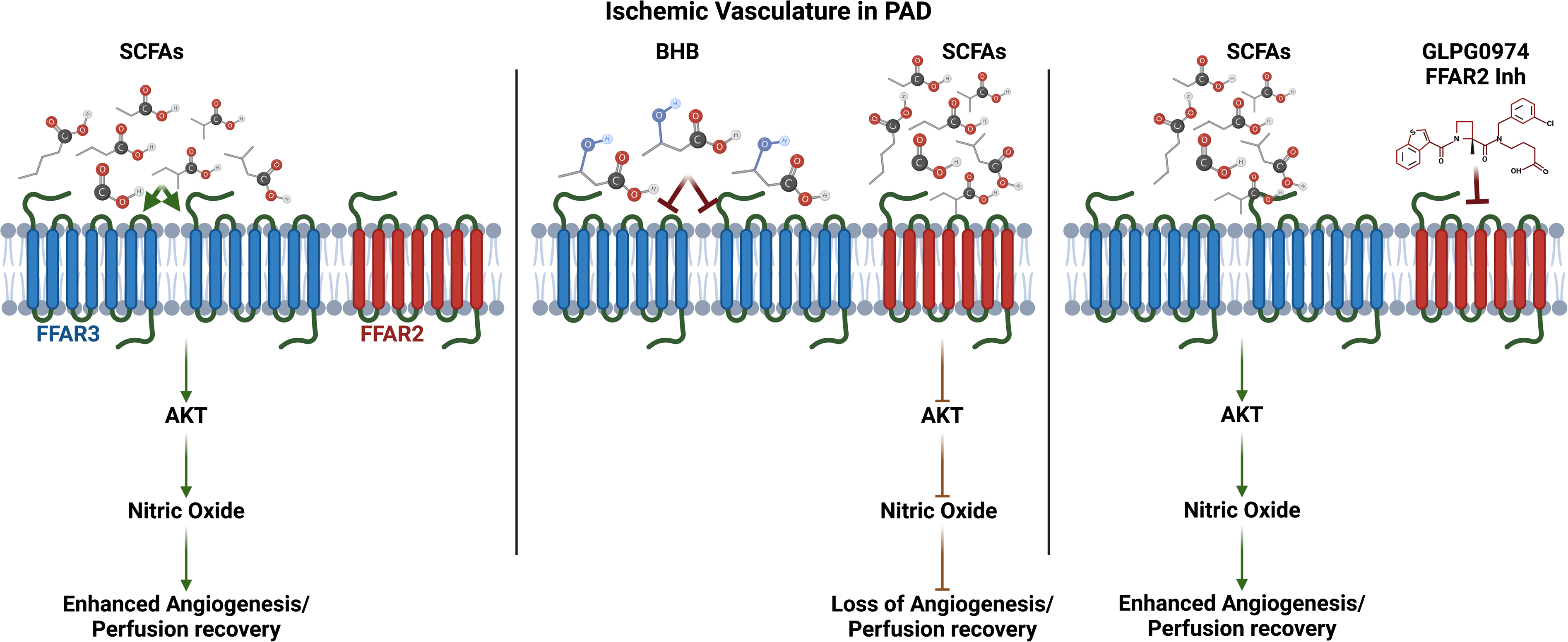
Graphical Abstract/schematic of the study: Ischemia induces FFAR3 expression on endothelial cells increasing their bioavailability for SCFAs to bind and induce angiogenesis via the AKT-NO pathway (left panel). Inhibiting FFAR3 by BHB inhibits the AKT-NO pathway thereby decreasing the angiogenic capacity of ischemic-ECs (middle panel). inhibiting FFAR2 by GLPG0974 facilitates SCFAs to bind and activate FFAR3 to induce ischemic-angiogenesis (right panel).

Next, to determine the causal role of FFAR2 in driving ischemic-EC angiogenic capacity, we treated ischemic-ECs with FFAR2-inhibitor dose-dependently in the presence or absence of SCFAs. *In vitro* angiogenesis assay showed that inhibiting FFAR2 significantly induced the angiogenic capacity of ischemic-EC in a dose-dependent manner vs. controls (%No of Nodes: 10µM-P<0.0001, %No of Junctions: 10µM-P=0.028, %No of Branches: 5µM-P=0.01, 10µM-P<0.0001, Fig-7C). SCFAs induced the angiogenic capacity of ischemic-ECs treated with FFAR2 inhibitor at all concentrations (Fig-7C). This data indicated that FFAR2 negatively controls the angiogenic capacity of ischemic-ECs; and the ability of SCFAs to induce ischemic-EC angiogenic capacity is not dependent on FFAR2. However, inhibiting FFAR3 by BHB treatment (Fig-7D) or silencing FFAR3 (Fig-7E) significantly blunted the pro-angiogenic response observed in FFAR2-inhibited ischemic-ECs. Furthermore, SCFAs failed to induce ischemic-EC angiogenic capacity in FFAR2-inhibited ischemic-ECs when treated with BHB (Fig-7D) or FFAR3 is silenced (Fig-7E) indicating that the ability of SCFAs to induce ischemic-angiogenesis is dependent on the dominant pro-angiogenic role of FFAR3 in ischemic-ECs (Fig-7D, E).

Silencing-FFAR3 or inhibiting-FFAR3 by BHB significantly decreased maximal-respiration but did not significantly affect ATP-levels (Fig-6C, F). Moreover, SCFAs further decreased maximal-respiration in FFAR3-silenced or FFAR3-inhibited ischemic-ECs, but did not affect ATP-production (Fig-6C, F). This data indicates that the changes observed in the angiogenic-capacity of ischemic-ECs treated with SCFAs with or without FFAR3-inhibition are independent of ATP-production^48^. Since SCFAs induce AKT-activation that is upstream of NO-production^49^, we wanted to determine whether SCFA-FFAR3/FFAR2 signaling controls NO-production to regulate ischemic-EC angiogenic capacity. Consistent with the ability of SCFAs to induce AKT-activation, ischemic-ECs treated with SCFAs showed a significant increase in NO levels vs. controls (P=0.018, Fig-7F; P<0.0001, Fig-7G). While FFAR2-inhibition, FFAR3-silencing, or FFAR3-inhibition did not affect NO-production by themselves (Fig-7F, G); blocking FFAR3 function inhibition blocked SCFA-induced NO-production in ischemic-ECs. FFAR2-inhibition rescued the loss of NO-production capacity by SCFAs in FFAR3-inhibited ischemic-ECs (Fig-7F, G). This data indicated that FFAR3-inhibition blocks the NO-production capacity of SCFAs in ischemic-ECs. SCFAs were well known to control inflammation by decreasing ROS production^50,51^. Hence, we next examined whether the loss of perfusion recovery in BHB+SCFA-treated ischemic-muscle is due to changes in redox balance. H2DCFDA, a cell-permeant indicator or reactive oxygen species (ROS)^52^ showed that while SCFAs, FFAR2-inhibition, or FFAR3-inhibition did not affect ROS levels (Fig-7H, I), SCFAs significantly induced ROS levels in FFAR3-inhibited ischemic-ECs. FFAR2-inhibition blocked the ROS levels in FFAR3-inhibited ischemic-ECs treated with SCFAs. This data indicated that the mechanism by which BHB+SCFA treatment impairs perfusion recovery and inhibits ischemic-EC angiogenic capacity is by decreasing NO and increasing ROS levels thereby inhibiting the redox balance in ischemic-ECs.

## Discussion

Detrimental effects of Palmitate have been shown in multiple cardiovascular pathologies and EC functional impairment^13,53,54^. *In vitro,* studies have shown that Palmitate induces inflammatory cytokine production including TNF-α, IL-6, and MCP1, LDL1 in macrophages and decreases NO-production in ECs^14,55–57^. More importantly, Palmitate has been associated with atherosclerotic plaque vulnerability and major cardiovascular adverse events in T2D^58,59^. However, whether increased Palmitate levels can be the sole determinant of EC dysfunction and impaired perfusion recovery in PAD is not known. Consistent with our previous study, Palmitate significantly decreased ischemic-EC survival and angiogenic capacity^36^ dose-dependently but did not affect the EC barrier. Accordingly, we anticipated that the delivery of Palmitate into the ischemic-muscle would recapitulate the severe loss of perfusion recovery and significantly higher necrosis scores/severity observed in type-2 diabetic PAD models (T2D-PAD, C57BL/6J mice on a high-fat diet (60kCal) for 3-4 months)^36,38^. However, the magnitude of perfusion loss did not match the severity observed in T2D-PAD mice. This data indicated that while increased Palmitate levels in T2D-PAD could contribute to impaired perfusion recovery, Palmitate alone is not the driver of disease severity observed in T2D-PAD. Consistently, previous studies have shown that while 300mM Palmitate delivered intraperitoneally aggravated nonalcoholic fatty liver disease^60^, and 9nmol Palmitate delivered intracerebroventricular induced memory deficit ^61^; 15µM dietary Palmitate inhibited arthritis^62^. These data point towards that the concentration of exogenously delivered Palmitate directs the disease outcomes. In our current study, we used 250µM Palmitate i.m. matching with the minimal dose that decreased ischemic-EC survival by 50% in our *in vitro* experiments. Based on our data and these previous studies, while higher Palmitate concentrations could increase the disease severity in experimental-PAD, the pathological relevance of those concentrations will be unclear.

While Palmitate showed a detrimental effect on ischemic-EC survival and angiogenic capacity *in vitro*, LCFA treatment did not induce any significant changes. One of the reasons could be due to the lower individual LCFA levels in the 5% LCFA concentration. LCFA treatment at higher than 5% concentration resulted in loss of solubility in the culture medium limiting our studies to 5% LCFA treatment as a comparator. Further studies are needed to establish a suitable vehicle that can increase the solubility of LCFAs (individually or as a mix) in culture media to allow concentrations that match with Palmitate treatment for *in vitro* functional studies. Furthermore, previous studies have indicated distinct effects of LCFAs on ECs, e.g., Palmitate but not Palmitoleic induces insulin resistance^63^; Palmitate but not Stearic acid inhibits NO-production^64^; Palmitate inhibits whereas Oleate promotes EC proliferation^65^. This data indicates that distinct functions of LCFAs; and when combined can have a more complex or net neutral phenotypic response as observed in our LCFA-treated ischemic-ECs. In contrast with Palmitate, SCFAs did not show any significant effect on normal-EC proliferation or ischemic-EC survival. However, SCFAs showed a significant increase in the angiogenic capacity of both normal and ischemic-ECs vs. controls. Consistently, SCFAs significantly induced AKT activation and downstream NO-production in ischemic-ECs Furthermore, while SCFA was able to improve the normal-EC barrier, no significant effect on the ischemic-EC barrier was observed. This data indicated that the primary effect of SCFAs on ischemic-ECs was in regulating their angiogenic capacity.

Since, LCFAs and Palmitate undergo β-oxidation in the mitochondria by the Carnitine shuttle system^31^, and ECs depend on glycolysis^42^ primarily for metabolic requirements, we determined the metabolic processes regulated by Palmitate and SCFAs in controlling ischemic-EC function. Seahorse metabolic assays showed that Palmitate did not affect normal-EC glycolysis, respiration, or basal-FAO. However, in the ischemic-ECs, Palmitate significantly decreased glycolysis, respiration, and basal-FAO. Paradoxically, despite a decrease in glycolysis and respiration, Palmitate significantly induced PFKFB3 (glycolysis) and SDHA (TCA-cycle) levels indicating a compensatory metabolic outcome. It is important to note that SCFA-induced ischemic angiogenesis occurred by decreasing glycolysis in ischemic-ECs. This data was consistent with our recent reports showing that blocking ischemia-induced maladaptive glycolysis promotes ischemic-EC angiogenic capacity^40,52,66^. Furthermore, no significant difference in the ATP-production in ischemic-ECs treated with SCFAs indicated that the ability of SCFAs to drive ischemic angiogenesis is not dependent on driving mitochondrial ATP production. Even though both FFAR2 and FFAR3 are primary receptors for SCFAs ^20^, in the ischemic-ECs, SCFAs showed a preference for FFAR3 over FFAR2. This bias of SCFAs towards FFAR3 is reflected in the pro-angiogenic effects of FFAR3 vs. anti-angiogenic effects of FFAR2, since silencing FFAR3 inhibited ischemic-EC angiogenic capacity, whereas inhibiting FFAR2 induced ischemic-angiogenesis. Furthermore, contrary to our prediction, LC-MRM MS analysis showed no significant difference in SCFA levels in ischemic-muscle vs. non-ischemic. In fact, a significant decrease in the levels of Dodeconic acid levels and a significant increase in FFAR3 levels was observed in the ischemic-muscle vs. non-ischemic indicating that lower SCFA levels in the ischemic-muscle is a limiting factor in promoting perfusion recovery in PAD. While pharmacological inhibitors for FFAR2 are available^67,68^, the only known pharmacological inhibitor for FFAR3 is BHB^26,27,29,30^. Despite the relative similarities in their chemical structure, Butyrate and BHB were shown to exert distinct functions. E.g. Chriett et al^27^., indicate that Butyrate but not BHB is a strong HDAC1-3 inhibitor in myotubes. Consistent with this data, we did not observe any significant effect of BHB on HDAC1-3 expression but a significant decrease in FFAR3 expression in ischemic-ECs.

In our efforts to confirm the causal role of FFAR3 in driving SCFA-induced perfusion recovery, we inadvertently, noticed that inhibiting FFAR3 via BHB treatment shifted the pro-angiogenic SCFA function to anti-angiogenic and impaired perfusion recovery. It is important to note that SCFA treatment in FFAR3-inhibited C57BL/6J mice showed a numerical increase in necrosis incidence, a mice strain that recovers better from HLI with limited tissue loss in ischemic-muscle^45,69^. Mechanistically, our data showed that the loss of FFAR3 increases the bio-availability of SCFAs to FFAR2 resulting in decreased NO-production capacity of SCFAs; and increased oxidative stress in ischemic-ECs to inhibit ischemic-EC angiogenic capacity.

## Conclusions

Severe PAD often results in progressive limb amputation with an accompanying 5-year mortality rate of ∼35%^70^. Currently, no known therapies can directly address/correct the impaired blood flow and perfusion to the ischemic limb and provide perfusion relief to patients with PAD^37^. Hence, targeting SCFAs/gut microbiome by adjusting the diet of patients with PAD is an attractive and feasible therapeutic approach. However, the severity of vascular occlusions that limit the blood supply to the ischemic-muscle of patients with PAD can also decrease the delivery of circulating SCFAs to the distal ischemic-muscle thereby limiting a therapeutic response. Furthermore, SCFA treatment decreased perfusion recovery upon FFAR3 inhibition by BHB. This data points towards the necessity for personalized/precision medicine for PAD since SCFA treatment or targeting gut microbiome to increase SCFA levels in patients with PAD with lower FFAR3 levels in ischemic-muscle can result in further impairment.

## Gaps and Future directions

Currently, the relative or actual levels of SCFAs (or LCFAs) in patients with PAD patients is not known. More importantly, sex as a determinant in regulating SCFA-induced perfusion recovery needs to be further investigated. Furthermore, ischemia did not significantly change the overall SCFA content in the skeletal muscle suggesting that a loss of SCL16A family members in ischemic-muscle might limit their uptake from circulation thereby contributing to decreased perfusion recovery. While BHB inhibited FFAR3 levels, BHB as a ketone body can have functions beyond FFAR3 inhibition. Hence, future studies using EC-specific FFAR3-KO/FFAR2-KO mice in preclinical-PAD will help to determine the functional significance of these fatty acid receptors in controlling PAD outcomes.

## Funding

This work was supported by 7R01HL146673-02 to V.C.G.

## Disclosures

An abstract of this study was submitted for presentation at the AHA-Vascular Discovery conference-2025. The authors declare no conflict of interest.

## Author Contributions

SK, KP, and WZ contributed to the experiments, analyzed the data, and edited the manuscript. VG conceived, and supervised the study; analyzed the data; drafted, and edited the manuscript; and obtained funding for the study.

## Notes

### Competing Interest Statement

The authors have declared no competing interest.

